# Unmasking the large and highly repetitive genome of *Phlox* reveals a complex evolutionary history of speciation with gene flow

**DOI:** 10.64898/2026.09.01.748350

**Authors:** Felix L. Wu, Danielle E. Khost, Patrick F. McKenzie, Samridhi Chaturvedi, Grace A. Burgin, Timothy B. Sackton, Robin Hopkins

## Abstract

The large size and high repetitive content of many plant genomes have hindered elucidation of how the fundamental processes of evolution give rise to speciation. Here we present chromosome-level assemblies of 6 gigabase genomes for four closely related *Phlox* wildflower species. We generate extensive population genetic data for the three well-studied annual species *P. drummondii, P. cuspidata*, and *P. roemeriana*, including structural variation from whole-genome long-read resequencing data. The whole-genome assemblies reveal extensive differences in amount and distribution of genetic variation within and between species, reflective of differences in life-history strategies, mating systems, and edaphic specialization. The population genetic data exposes a history of widespread and consistent gene flow between all three annual *Phlox* species throughout their divergence and speciation. Our unmasking of structural variants and repetitive elements exposes rich and dynamic forms of genetic variation that show strong phylogenetic signal and pervasive patterns of gene flow. Our results lay a foundation for untangling the highly complex genomes of plants to make advancements in our understanding of the processes of speciation and divergence.

**Significance:** Plant genomes are often very large and filled with repetitive DNA which has made using genomics to study plant evolutionary dynamics and the formation of new species difficult. To address these challenges, we produced high-quality genome assemblies for four closely related wildflower species in the genus *Phlox*. By comparing genetic variation across these species, we show that genetic exchange has persisted throughout their evolutionary split. We demonstrate how this history can be made visible in messy, repetitive DNA. Genomic variation among the species also reflects differences in their reproductive strategy and habitat. These findings show that that these wildflowers have porous species boundaries and provide a valuable toolkit for studying other complex plant genomes.

## Introduction

Flowering plants have long held distinct historical importance for understanding the evolutionary processes of speciation and hybridization^1^. Three centuries of work by botanists, naturalists, and agronomists have laid a foundation for our understanding of selection during the process of divergence, the impacts of gene flow on diversification, and the evolutionary importance of hybridization^2–4^. Ready natural observation, feasibly controlled breeding, and ease of manipulation in common gardens experiments have made plants ideal for inferring how fundamental mechanisms of evolution shape the process of speciation^5,6^. Now, modern techniques for analyzing genetic variation have further expanded our ability to detect genome-wide signatures of selection, gene flow, divergence, and the consequences of hybridization, yet applying these methods to plants has proven challenging^7–10^.

Angiosperms have notoriously flexible genomes that vary over 24,000-fold in size^11^. While hybridization and whole genome duplication (polyploidization) are obvious and abrupt mechanisms of evolving genome complexity, increasing attention has been paid towards the importance of selfish transposable elements (TEs) in shaping genome size and genetic variation in flowering plants^12,13^. TEs can generate enormous tracts of repetitive sequences that vary in size and location between and even within species^14,15^. The abundant genetic variation associated with TEs is largely structural in the form of insertions and deletions and has made genome assembly and analysis difficult for many organisms and especially plants^16^. Methods that infer patterns of evolution from genomic data typically involve masking of repetitive and structural variation^17–19^. Ignoring such “messy” genetic variation could hide signatures of key evolutionary processes. After all, TE activity has repeatedly been implicated in speciation by causing reproductive isolation through hybrid dysregulation and generating mutations leading to adaptive trait variation and divergence^20–23^. Therefore, unmasking this complex genetic variation is critical for a complete understanding of whole genome variation and can lead to stronger inferences about fundamental evolutionary processes such as selection and gene flow during speciation.

The wildflower genus of *Phlox* (*Polemoniaceae*) is an ideal system to investigate complex genetic variation and evaluate its importance for evolutionary inference. Historically, *Phlox* has acted as a critical plant system for understanding fundamental evolutionary processes including the costs and benefits of mating system variation, distribution and dispersal of adaptive traits, and co-evolution and pollination biology^24–30^. More recently, it has developed into an important model for studying speciation. Hybridization plays a key role in diversification of this group through the formation of allopolyploid species, and gene flow is evident between an abundance of sympatric species^31,32^. Particular attention has been given to a small clade of three annual species—*P. drummondii, P. roemeriana*, and *P. cuspidata*—that harbor the best characterized case-study of reinforcement in which natural selection drives the evolution of prezygotic reproductive isolation to prevent costly hybridization^26,27^. *P. drummondii* occupies a native range in central and eastern Texas, USA, with *P. cuspidata* and *P. roemeriana* inhabiting overlapping ranges to the east and west respectively. In the region of sympatry with *P. cuspidata, P. drummondii* evolved a dramatic shift in flower color from the ancestral light blue color common to most *Phlox*, to dark red. The derived dark red flower color decreases hybridization between the sympatric species by 50% relative to the ancestral light blue form^26^. This textbook example of reinforcement suggests an evolutionary history of gene flow across these species, yet the timing and nature of such events—much less the latent pattern of genomic structure and organization within each species—remain unknown. Equally uncertain is the extent to which these patterns are reflected across the entire genome: whether TE sequences and well-behaved compartments of the genome experience the same fate after introgression and divergence.

Genomes of various *Phlox* species have been karyotyped, and are estimated to have large genomes (> 6 Gb) packaged onto few enormous chromosomes (1*n* = 7)^33–35^. The size and complexity of the *Phlox* genome have previously been barriers to whole genome inference methods. Here we demonstrate that unmasking this enormous wealth of genetic diversity can provide new insights into the speciation process.

We present whole genome assemblies for each species of the monophyletic clade of annual *Phlox*, as well as the outgroup perennial species, *P. pilosa*. We complement these genomes with extensive population genetic and phylogenomic resequencing data using both short and long-read technologies. We leverage the genomes of these species to infer the evolutionary history of divergence and diversification with special attention to how hybridization and gene flow have shaped the patterns of speciation in this group. We assess the complex variation associated with repetitive regions and structural variants (SVs) through analyses of the distribution of TEs and their contributions to the genomic landscape. We disentangle a history of gene flow within the clade and demonstrate how to reveal the evolutionary signals of divergence and gene flow in unmasked, repetitive, structural variation. In so doing, we connect the rich history of natural observations and organismal studies to patterns of genomic variation across the species.

## Results and Discussion

### Assembly and annotation of the *Phlox* genome

To resolve patterns of divergence and speciation in *Phlox*, we sequenced and assembled the genome of a *Phlox drummondii* individual (Fig. 1A). We generated a *de novo* assembly using 241 Gb of Oxford Nanopore Technologies (ONT) long-reads, equivalent to 35x coverage. We scaffolded the initial 28,022 contigs using both a Bionano optical genomic map and a genetic linkage map. The final haploid assembly contains 6.3 Gb of total sequence (7.4 Gb gapped) and is highly contiguous (scaffold/contig N50 = 406 Mb / 668 kb) across seven massive chromosomes ranging in size from 375 to 854 Mb (Table 1). Genome size is consistent with previous size estimates obtained from flow cytometry (5.98 Gb)^35^. Benchmarking with universal single-copy orthologs indicates that our genome is highly complete (BUSCO > 99%) (Table 1).

**Table 1.**
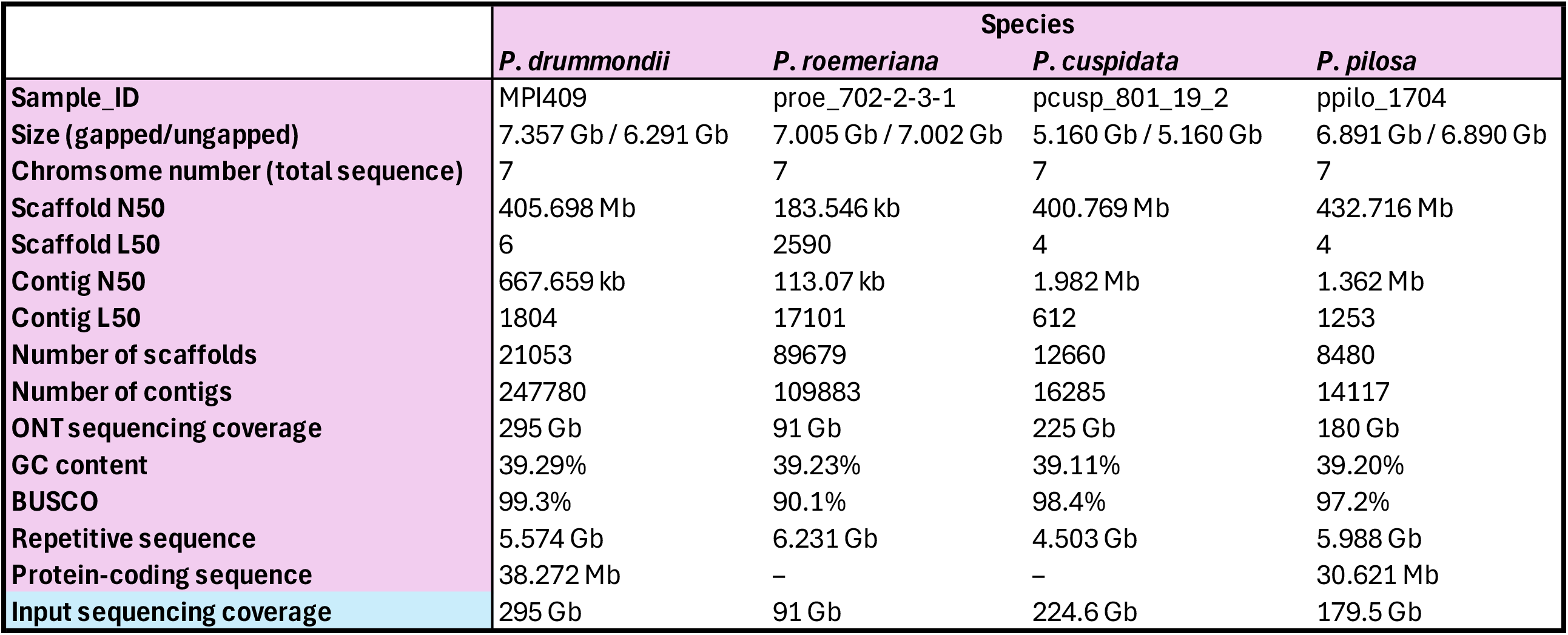
Genome assembly statistics.

**Figure 1.**
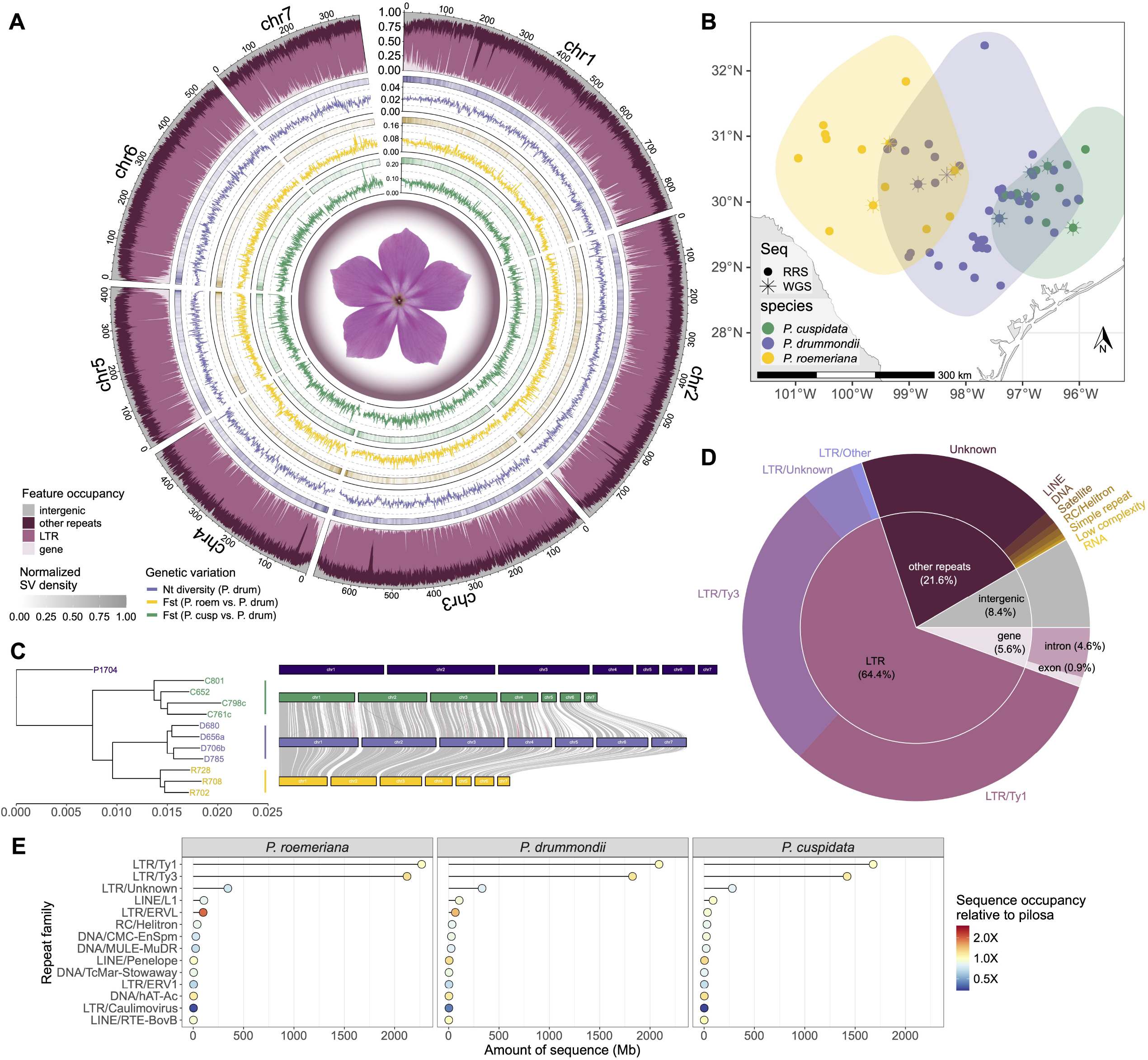
Genome assemblies and content of annual *Phlox*. (A) Circular plot of assembled chromosomes of *P. drummondii* genome. Outer track displays proportions of 1 Mb genomic windows occupied by genic (light pink), long terminal repeat (LTR, dark pink), non-LTR repeat (burgundy), and intergenic (gray) features. Purple lines along the second outermost track denote nucleotide diversity (*π*) estimated across four *P. drummondii* individuals in 1 Mb windows across the genome. Yellow and green lines of the two innermost tracks denote *F*_*ST*_ estimates in 1 Mb windows between *P. drummondii* and *P. roemeriana* and between *P. drummondii* and *P. cuspidata*, respectively. Shading in narrow bands above each line plot indicates—for each window—the density of structural variants identified in long-read sequences from *P. drummondii* (purple), *P. roemeriana* (yellow), and *P. cuspidata* (green). (B) Geographic distribution of sampled annual *Phlox* species across the state of Texas, USA. Points denote population sampling locations of *P. roemeriana* (yellow), *P. drummondii* (purple), and *P. cuspidata* (green). Circles correspond to samples subjected to reduced-representation sequencing (RRS) while stars denote sampling locations for whole genome sequencing (WGS). Shaded areas indicate approximate ranges of each species. (C) Phylogeny and syntenic comparison of assembled *Phlox* species. Left portion shows inferred phylogenetic relationships between species using WGS samples. Right portion shows relative sizes of assembled chromosome of *P. pilosa* (dark purple), *P. cuspidata* (green), *P. drummondii* (purple), and *P. roemeriana* (yellow) top to bottom. Gray bands denote syntenic blocks between *P. drummondii* and the other two annual species while red bands indicate inversions. (D) Summary of feature occupancy across the *P. drummondii* genome. Inner pie chart is segmented into the same categories as in the outer track of (A); outer ring subdivides repeats into families. Sector sizes correspond to proportion of base pairs occupied. (E) Lollipop plots of repetitive element sequence occupancy in three Texas annual *Phlox*. Positions of each point along x-axis denote absolute base pair occupancy of a particular repeat element family (y-axis). Points are colored according to whether the proportion of genome occupied by a given element family is greater than (red) or less (blue) than that of outgroup species *P. pilosa*.

We annotated 35,223 genes—composing roughly 0.9% of the genome—using a transcriptome derived from RNA-Seq libraries sequenced from multiple tissue sources. Nucleotide diversity (*π*) is up to 50% reduced in exonic regions relative to the rest of the genome (Fig. S1; bootstrap *p* < 0.001), as measured using single nucleotide polymorphisms (SNPs) inferred from four independently sequenced *P. drummondii* individuals (Fig. 1A, see next section); these results are consistent with our expectation that negative selection operates more strongly on functional elements.

To infer broader evolutionary patterns across the species group we long-read sequenced and assembled whole genomes for the remaining annual species *P. roemeriana* and *P. cuspidata* as well as the outgroup perennial species *P. pilosa* (Fig. 1B). We achieved high levels of contiguity by scaffolding these genomes against the *P. drummondii* assembly (Table 1). Pairwise alignment of our four *Phlox* genome assemblies with Anchorwave^36^ reveal highly collinear chromosomes with a handful of relatively small inversions and translocations (Fig 1C).

In *P. drummondii*, repetitive sequences comprise 86% of the genome (Fig. 1D)—an especially large fraction on par with other large plant genomes such as maize (*Zea mays*, 83%)^37^ and wheat (*Triticum aestivum*, 90%)^38^. In *Phlox*, as in most plants, these repetitive sequences are predominantly long terminal repeat (LTR) retrotransposons (64.4% of the genome) (Fig. 1D). Local repeat content (base pair proportion) is reduced within annotated exons (bootstrap *p* < 0.001), and yet abundant in introns and gene-adjacent regions of the genome (Fig. S2). The consolidation of genes and TEs together at the ends of chromosomes in *Phlox* is consistent with the hypothesis that TEs (and particularly LTR retrotransposons) are involved in gene regulation and generating diversity in expression and isoform production^39,40^. Categorization of the abundance of distinct TE superfamilies^41^ revealed species-level differences in TE occupancy across the genomes. The perennial species *P. pilosa* harbors more copies of Caulimovirus, a close relative of LTRs^42^, compared to the three annual species, and endogenous retroviral activity has increased in the *P. roemeriana-drummondii* species pair relative to the other *Phlox* species (Fig. 1E).

### Comparative genomics reflects variation in life history strategy

We implemented a two-pronged comparative population genomics approach in *Phlox* that allows for inferences based on both breadth and depth of coverage. We used reduced-representation sequencing (RRS) to characterize 779 individuals from across 89 populations of the three Texas annual *Phlox* as well as 13 individuals of perennial out-group *P. pilosa* (Fig. 1B). We complemented this sampling with whole genome ONT long-read sequencing (WGS) of three *P. roemeriana*, four *P. cuspidata*, one *P. pilosa*, and four additional *drummondii* individuals (Table S1). Together, these datasets capture variation across both geographic (RRS) and genomic space (WGS) of this *Phlox* clade.

Phylogenetic tree reconstruction using WGS data yields a species topology consistent with previous reports^31,32^ showing *P. drummondii* and *P. roemeriana* are sister species to the exclusion of *P. cuspidata* (Fig. 1C). Annual *Phlox* samples in the RRS dataset likewise cluster by species when subjected to principal component analysis (PCA) and ancestry modeling with ADMIXTURE^43^. The first two PCs explain 27.3% and 12.1% of the genetic variation, respectively, and much of the improvement in cross-validation error was achieved with *K*= 3 populations (Fig. S3). These analyses also reveal strong evidence of admixed individuals that contain genetic variation associated with multiple annual *Phlox* species (Fig. 2A–B). The genetic divergence inferred between *Phlox* species varied depending on sequencing methods, with somewhat lower divergence of the annuals from *P. pilosa* for the RRS data (median *d*_*XY*_ = 0.91%) compared to the WGS data (median *d*_*XY*_ = 2.94%). Whole genome sequencing reveals moderate divergence between *P. drummondii* and the other two annual *Phlox* species with higher *F*_*ST*_ with *P. cuspidata* (*F*_*ST*_ = 0.146) than with *P. roemeriana* (*F*_*ST*_ = 0.104). Scans across the genome using 1 Mb windows reveal regions of exceptional divergence showing loci with consistently high *F*_*ST*_ against both *P. cuspidata* and *P. roemeriana* and low diversity within *P. drummondii* (such as the end of Chromosome 2 and the end of Chromosome 7) (Fig 1A, Fig. S4). These patterns motivate future inquiries into signals of selective sweeps and adaptive divergence within the *P. drummondii* species.

**Figure 2.**
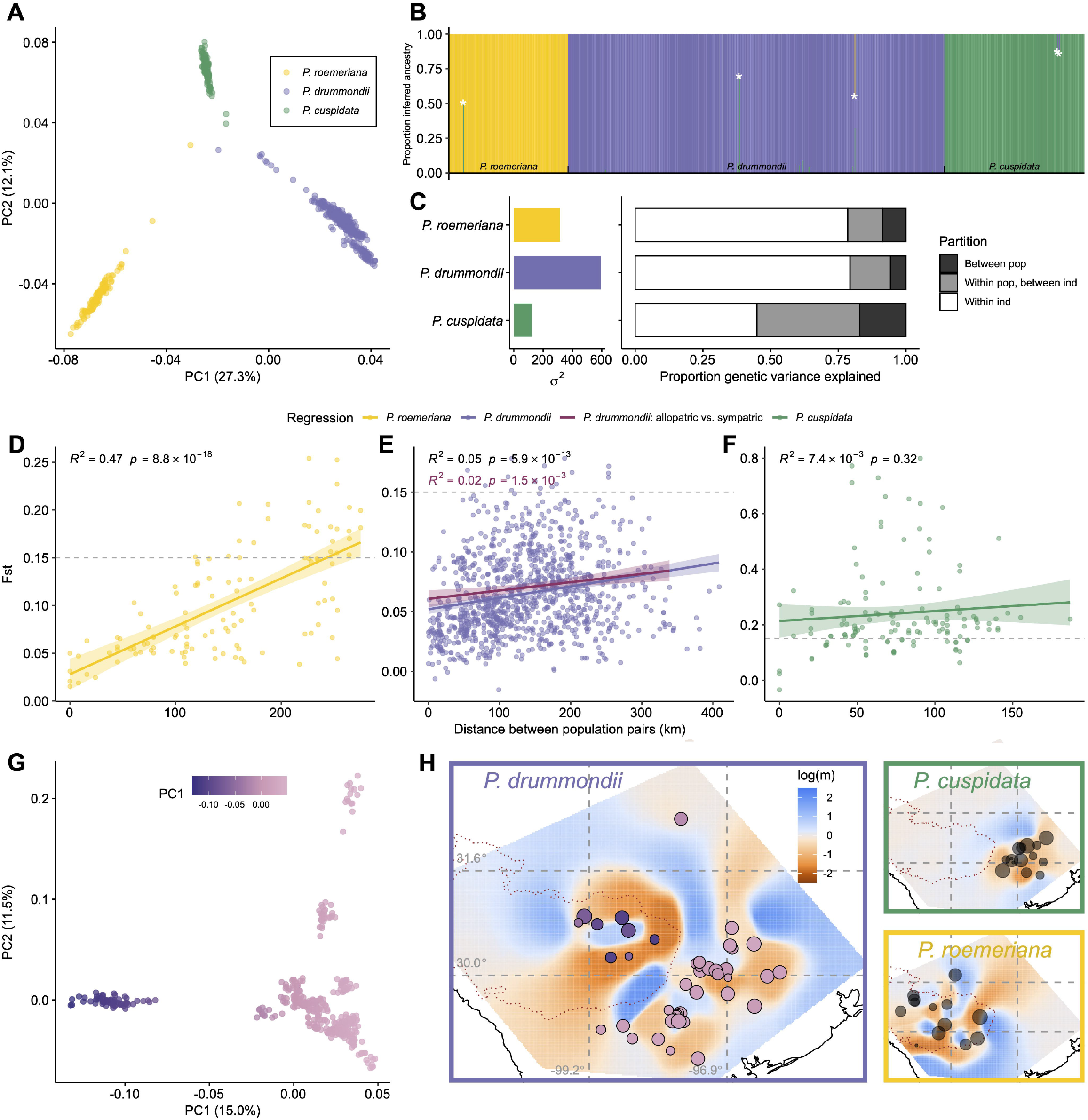
Patterns of genetic variation in annual *Phlox*. (A) Principal components analysis (PCA) of *P. roemeriana, P. drummondii*, and *P. cuspidata* individuals using 9,536 genotyped markers from reduced-representation sequencing (RRS). Individuals are colored by species labels as determined from phenotypic appearance in the field. (B) Mixing proportions for *K*= 3 ancestral populations determined with ADMIXTURE using the RRS dataset. Heights of colored bars denote estimated ancestry proportions corresponding to each of the three populations. Individuals are grouped by species label (black ticks) and within each group arrayed from left to right by increasing longitude. White asterisks denote individuals exhibiting recent admixture, i.e., with a minimum ancestry proportion across all populations > 0.9. ADMIXTURE plots for larger values of *K* are available as Supplemental Fig. S9. (C) Analysis of molecular variance (AMOVA) of each *Phlox* species. The left subplot shows absolute levels of genetic variance, *σ*^2^, for species while the right subplot decomposes the relative proportions of variance explained by within-individual (white), within-population (gray), and between-population (black) effects. Isolation by distance for *P. roemeriana* (D), *P. drummondii* (E), and *P. cuspidata* (F). Each point denotes a pair of sampled populations and their corresponding *F*_*ST*_ value (y-axis) and inter-population distance in kilometers (x-axis). Lines and shaded bands indicate ordinary least squares regression fits and 95% confidence intervals, respectively; model fit parameters are given in the upper left corner of each figure. Grey dotted lines are drawn at *F*_*ST*_ = 0.15 to aid visual comparison between species. For *P. drummondii* (E), the purple line and text denote regression using all points while the dark pink line and text denote regression using only population pairs in which one *P. drummondii* population is sympatric with *P. cuspidata* and the other is allopatric. (G) PCA of genetic variation in *P. drummondii* individuals using 16,144 genotyped markers from the RRS dataset. Points are colored by PC1 value. (H) Estimated Effective Migration Surface (EEMS) plots for *P. drummondii* (purple box), *P. roemeriana* (yellow box), and *P. cuspidata* (green box). Borders of Texas are drawn in black, red dotted lines outline the Edwards Plateau ecoregion, and log-scaled relative migration rate estimates are colored from low (orange) to high (blue). Circular points denote locations of sampled populations in the RRS dataset and are sized relative to the number of individuals sampled from each. For *P. drummondii*, point colors correspond to mean PC1 value (from G).

Our RRS dataset is poised to characterize the distribution of genetic variation within individuals, population, and species. Our expectation for how genetic variation is distributed within this clade stems from extensive natural history knowledge of the mating systems and habits of the annual *Phlox* species. *P. drummondii* and *P. roemeriana* exhibit genetic self-incompatibility, necessitating outcrossing, whereas *P. cuspidata* is self-compatible with reportedly high selfing rates^44,45^. *P. drummondii* is the most widespread in its geographic range of the three annual species. *P. roemeriana* is more restricted, with small populations on the limestone outcrops of the Edwards Plateau in central Texas, USA, and *P. cuspidata* tends to grow in large, isolated populations throughout eastern Texas, USA (Fig. 1B)^29^. Based on these differences, we expect *P. drummondii* to harbor extensive genetic variation with limited population structure, while *P. roemeriana* and *P. cuspidata* will have higher population structure. Early work with allozyme data provides some support for this prediction^44^, though these initial results are limited in scope—drawing from only a handful of markers and locations.

As predicted, our analyses expose high inbreeding in *P. cuspidata* (median inbreeding coefficient = 0.66) and a reduction in its genetic variation relative to the other two annual species. The proportion of genetic variation explained by within-individual effects is similar in the two self-incompatible species but sharply reduced in *P. cuspidata* due to increased homozygosity from historical self-matings and biparental inbreeding (analysis of molecular variance [AMOVA] *p* < 0.002). Total genetic variance is two-to three-fold higher in *P. drummondii* compared to the other two annual *Phlox* species, consistent with this species being the most widespread, with the largest populations and high rates of outcrossing (Fig. 2C). Furthermore, the proportion of genetic variance explained by between-population differentiation is very low in *P. drummondii* (5.7%), and also strikingly consistent with earlier allozyme estimates (6%)^46^. The consistency in signal across different molecular marker type, number, and source individuals is a testament to the robustness of these fundamental population genetic parameters.

Each annual species exhibits a distinct pattern of genetic variation across the range. Isolation by distance (IBD) is strongest in *P. roemeriana* (*R*^2^ = 0.47, *p* = 8.8 × 10^−18^; Fig. 2D), achieving remarkably high *F*_*ST*_ at much shorter distances than the more widely distributed *P. drummondii* populations (*R*^2^ = 0.05, *p* = 5.9 × 10^−13^; Fig. 2E). This strong IBD in *P. roemeriana* is consistent with habitat specificity and isolation resulting in genetic isolation between populations^29^. While *F*_*ST*_ estimates between *P. cuspidata* populations are much higher than observed in the other two *Phlox* species, we find no pattern of IBD across the range of this species (*R*^2^ = 7.4 × 10^−3^,*p* = 0.32; Fig. 2F). A lack of IBD may arise if selfing rates are high relative to rates of seed dispersal^47^.

To investigate the evolutionary history of speciation in these species we begin by determining if reinforcement has left a population genetic signature across the geographic range of *P. drummondii*. Specifically, we compared *P. drummondii* populations that evolved dark red flowers to reduce costly hybridization in sympatry with *P. cuspidata*, to nearby allopatric *P. drummondii* populations that harbor the ancestral light-blue flower color. We found no evidence of genetic differentiation across the sharp phenotypic transition of flower color. Populations of different colors between allopatry and sympatry are not more genetically differentiated than is expected from background patterns of IBD (Fig. 2E; bootstrap test for greater intercept *p* = 0.994). This pattern indicates that the flower color variation responsible for restricting hybridization between species does not restrict gene flow between flower colors within the species, as is suggested by previous experimental data^27^. Instead, flower color differences between *P. drummondii* populations are maintained by strong selection in the face of extensive intraspecific gene flow.

*P. drummondii* does exhibit a distinct genetic cluster of individuals from the northeastern Edwards Plateau (Fig. 2G). We find relatively low levels of migration between the Edwards Plateau individuals and other *P. drummondii* populations as inferred by Estimation of Effective Migration Surfaces (EEMS; effective migration rate log(m) < −1.58 among bordering populations; Fig. 2H)^48^. This region of intraspecific isolation is within a range of sympatry with the sister species *P. roemeriana* in which we find parallel patterns of restricted migration with other *P. roemeriana* populations. These patterns of genetic structure within the three species suggests that intraspecific genetic differentiation is driven more by abiotic habitat factors that may similarly impact all the species, as opposed to biotic interactions between closely related species or speciation.

### Transposable element abundance reflects structural variation diversity and interspecies divergence

TEs act as both drivers and passengers of genome evolution, reshaping the genomic landscape through their selfish mobile nature while also subject to the wider forces of selection, drift, and gene flow acting on the organism within which they reside. TEs exhibit tremendous repetitiveness and high mutation rates^49–51^, and thus have historically been treated as genomic “dark matter” and ignored in evolutionary analysis and inference. However, long-read sequencing improves our ability to reliably identify TEs and their effects on genome structure, raising the possibility that TEs themselves may serve as useful evolutionary markers. Furthermore, quantifying TE activity can allow us to understand how TEs themselves evolve during speciation with gene flow.

Our first challenge in using TEs to infer the evolutionary history of our clade is defining the unit of variation that we can quantify and describe. Even small families of TEs can have hundreds of copies within a single genome, which themselves contain vast amounts of nucleotide variation. TE variation violates assumptions of standard phylogenetic and population genetic sequence analyses and thus makes inferring patterns of orthology and paralogy challenging^52^. Instead of using single nucleotide variation we use the patterns of insertions and deletions that TEs leave behind as they move around the genome. TEs generate mutations in the form of SVs through a variety of mechanisms, including by simple retrotransposition and by spawning repetitive sequences that increase the likelihood of chromosome misalignment during recombination, leading to indels and rearrangements^53,54^. Therefore, our first goal was to associate SVs with the TEs that caused the variation. We then characterize the abundance and patterns of these SVs between and within species to make inferences about species divergence and gene flow. SVs—like any mutation—are subject to an idiosyncratic combination of evolutionary forces, yet the close mechanistic relationship of TEs and SVs suggests their abundance patterns are coupled.

To identify SVs we aligned our ONT long-read WGS samples from all our *Phlox* species to the *P. pilosa* genome assembly, which minimizes reference bias. To reduce false positives, confirmation of an SV required agreement across three different SV calling algorithms. We limited our SV set to variants between 50 bp and 100 kb in size, ignoring unresolved breakpoints and translocations. We identified nearly 600,000 unique SVs (median 270,589 SVs per individual), with the vast majority characterized as deletions or insertions (56% and 44%, respectively). Most indels were less than 1 kb in length (interquartile range [IQR] = 86–477 bp), whereas duplications and inversions tended to be longer (IQR = 1508–6072 bp; Fig. S5). SVs were notably overrepresented in exons (1.3-fold enrichment; one-sided binomial test *p* « 0.001) and introns (3.3-fold enrichment; one-sided binomial test *p* «0.001), which occupy 0.9% and 4.6% of the genome, respectively (Fig. S6). We annotated each SV for associated TEs by passing the sequence tracts involved in the variants through our repeat masking pipeline. For each TE family, we then summarized the number of copies, *N*_*fam*_ found in a given set of SVs, *N*_*fam*_ [*SV*]. We also collated TE family copy number in the reference genome of each species, *N*_*fam*_ [*Ref*_*sp*_]. We term the value *N*_*fam*_ the “TE-SV count” (Fig. 3A). As we predicted, for each annual *Phlox* species we observe a high degree of correlation between genomic abundance of TE families (*N*_*fam*_ [*Ref*_*sp*_]) and TE-SV counts (*N*_*fam*_ [*SV*_*sp*_])—in other words, the most abundant TEs also generate the most SVs (Fig. 3B).

**Figure 3.**
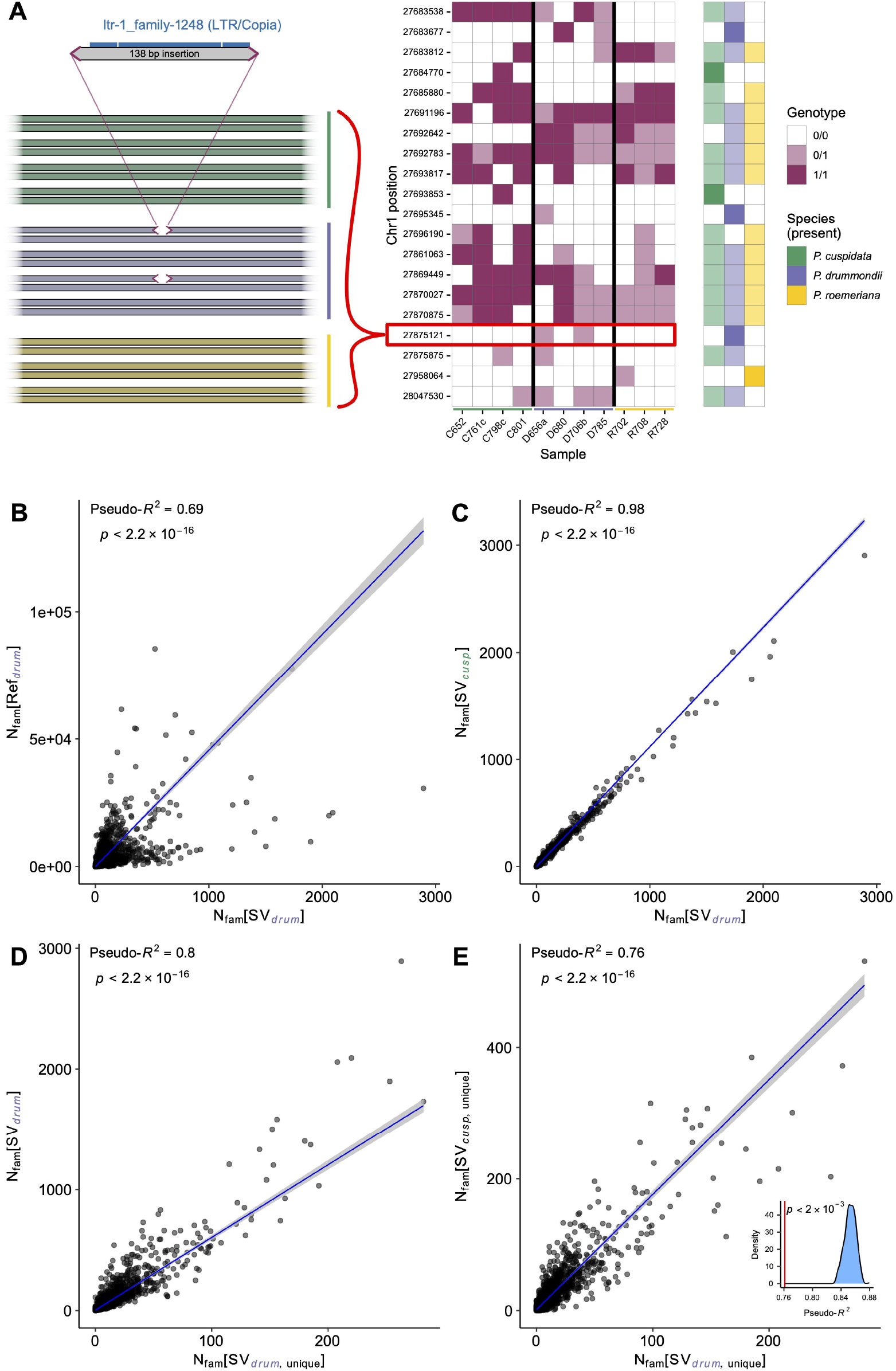
Transposable element dynamics and structural variation. (A) Schematic of procedure for annotating and associating structural variants (SVs) with transposable element (TE) families. First structural variants and associated transposable element were identified as shown on top-left by sequence (gray) from a 138 bp insertion (relative to the *P. pilosa* reference) carrying a Ty1-Copia long terminal repeat (LTR) element (blue) as annotated by RepeatMasker. Second, structural variation was genotyped across the entire whole genome sequencing (WGS) dataset shown here for the 138bp insertion present in heterozygous state in two *Phlox* genomes. Horizontal cartoon bars denote chromosomes for *P. cuspidata* (green), *P. drummondii* (purple), and *P. roemeriana* (yellow). Genotype data for all TE associated SVs were compiled for each individual and species; as an example, genotypes for twenty neighboring SVs on Chromosome 1 (*P. pilosa* reference) are displayed in a matrix (middle panel) with rows corresponding to SVs and columns corresponding to individual genomes. White, light pink, and dark pink cells indicate homozygous reference, heterozygous, and homozygous SV genotypes, respectively. Presence of a particular SV in each species is summarized by the shaded matrix to the right with darker cells indicating SVs unique to a single species. Red box denotes SV shown in left panel. (B-E) TE family copy numbers across different subsets of structural variants (SV) observed in annual *Phlox* species; each point denotes a distinct TE family, blue lines indicate negative binomial regression fits, and grey regions indicate 95% confidence intervals on the regression parameters. (B) TE family counts across *P. drummondii* reference genome versus counts across SVs present in *P. drummondii*. (C) TE family counts across SVs present in *P. cuspidata* vs SVs present in *P. drummondii*. (D) TE family counts across all SVs present in *P. drummondii* vs TE-SV counts unique to *P. drummondii*. (E) TE family counts across SVs unique to *P. cuspidata* vs TE-SV counts unique to *P. drummondii*. Inset plot shows bootstrap distribution of pseudo-*R*^2^correlations expected by chance in TE-SV counts unique to *P. cuspidata* vs *P. drummondii*, compared to the observed value, denoted by the red line.

Each species of annual *Phlox* was genotyped across multiple individuals for presence and absence of each SV. This dataset of SVs and associated TEs provides a window into the evolutionary histories of the TEs, since the relative age of SVs can be inferred from segregation patterns between and within species. For example, SVs that are polymorphic across all three annual *Phlox* species are inferred to be ancient mutations in the ancestor of the clade, whereas an SV polymorphic in just one species is likely due to a more recent movement of a TE in the specific lineage. We find broad correlation in the TE-SV counts of SVs polymorphic within all annual species and TE-SV counts of SVs observed in only one species (*p* < 2.2 × 10^−16^, *R*^2^> 0.74 in all comparisons), suggesting that TEs active in the annual *Phlox* ancestor tend to also be active at recent evolutionary timescales within each extant species (Fig. 3C–D). Indeed, focusing only on SVs polymorphic within a single species, we also find substantial—albeit attenuated—correlation between species in the abundance of SVs resulting from particular TE families. For example, we find a strong correlation between the TE-SV counts of SVs uniquely found in *P. drummondii* compared to the TE-SV counts of SVs unique to *P. cuspidata* (*p*< 2.2 × 10^−16^, *R*^2^= 0.76, Fig. 3E) despite each individual polymorphism being found in only one lineage. Such correlations suggest that the relative activity levels of most TE families are similar after recent species divergence in *Phlox*; while the significantly weaker correlation (Fig. 3E inset; bootstrap *p* < 0.002) of species-unique TE-SV counts is consistent with TE activity beginning to diverge across the clade. We note that, *P. drummondii*-specific TE-SV counts explain a larger proportion of variance in *P. roemeriana* (*R*^2^= 0.81) than in *P. cuspidate* (*R*^2^= 0.76). This tighter association between *drummondii* and *roemeriana* TE-SV counts is consistent with the hypothesis that activity of TEs diverges with time. We have thus demonstrated that genetic variation caused by TE mobility has a phylogenetic signal at this short timescale.

A growing interest in the evolution of TEs has been stymied by methodological and analytical challenges. While “jackpot” instances of TE activity generating new adaptive traits^55–59^ have been documented, the high mutation rate and repetitiveness of TEs has historically made robust genome-wide analysis challenging. By associating TEs to SVs, we gain a foothold into this problem. We have created a pipeline for associating TE activity with genomic variation to perform evolutionary and population genetic analyses directly on this variation without depending on linked single nucleotide variants. Our work demonstrates that TE families active in present-day species were likely the same ones active in the ancestor, with limited families undergoing recent expansion within their host species. Furthermore, by linking TEs to SVs, we detect a phylogenetic signal of decaying in similarity going backwards in time. Future analysis will continue to leverage this dataset to identify patterns of selection and TE suppression^60^.

### Genomic variation reveals extensive gene flow across in *Phlox*

Flower color variation in *P. drummondii* evolved through reinforcement. The evolution of a dark red flower color is selectively favored in regions of sympatry with *P. cuspidata* because it decreases the production of sterile hybrids through manipulation of pollinator movement between species^26^. Despite extensive characterization of the contemporary forces of selection favoring reproductive isolation, little is known about historical gene flow patterns across the annual clade that set the stage for the evolution of reproductive isolation in sympatry. Extensive theory has revealed the delicate balance between selection and gene flow necessary for reinforcement to successfully drive the evolution of reproductive isolation in sympatry^61–63^ and yet inferring these historical forces remains elusive^64^. Here we use genomic analyses to infer the extent and timing of gene flow throughout the speciation process.

Patterns of genomic variation reveal a history of widespread gene flow among the Texas annual *Phlox*. Using variation across the 12 WGS individuals, we computed F_4_-ratio admixture statistics between all possible subtrees of single individuals from each of the three annual species, designating the *P. pilosa* individual as the outgroup. To account for correlations across overlapping comparisons, we summarized these F_4_-ratio statistics using the f-branch statistic^65^, which reveals multiple signals of inter-species admixture (Fig. S7). *P. drummondii* individuals drawn from the sympatric range exhibit persistent introgression signals with *P*. cuspidata. These patterns of admixture between *P. drummondii* and *P. cuspidata* are consistent with observations of hybrid individuals in mixed *Phlox* populations (Fig. 2B). Surprisingly, there are also signals of admixture involving *P. roemeriana* and *P. cuspidata*, which have not been observed to grow sympatrically in recent history.

To further disentangle the history of gene flow represented in our sequenced genomes, we examined D-statistic (a.k.a. ABBA-BABA test) values computed across all interspecific comparisons of four individuals (48 total comparisons, Fig. 4A). D-statistic values were largely invariant depending on the specific *P. cuspidata* individual. In contrast, different *P. drummondii* and *P. roemeriana* individuals exhibit highly variable levels of allele sharing with *P. cuspidata*. This is particularly true in the case of *P. roemeriana* for which a single individual, R702, shows a strong signal of admixture (−0.08 < *D* < − 0.01) suggesting inconsistent, localized, or very recent incidences of gene flow.

**Figure 4.**
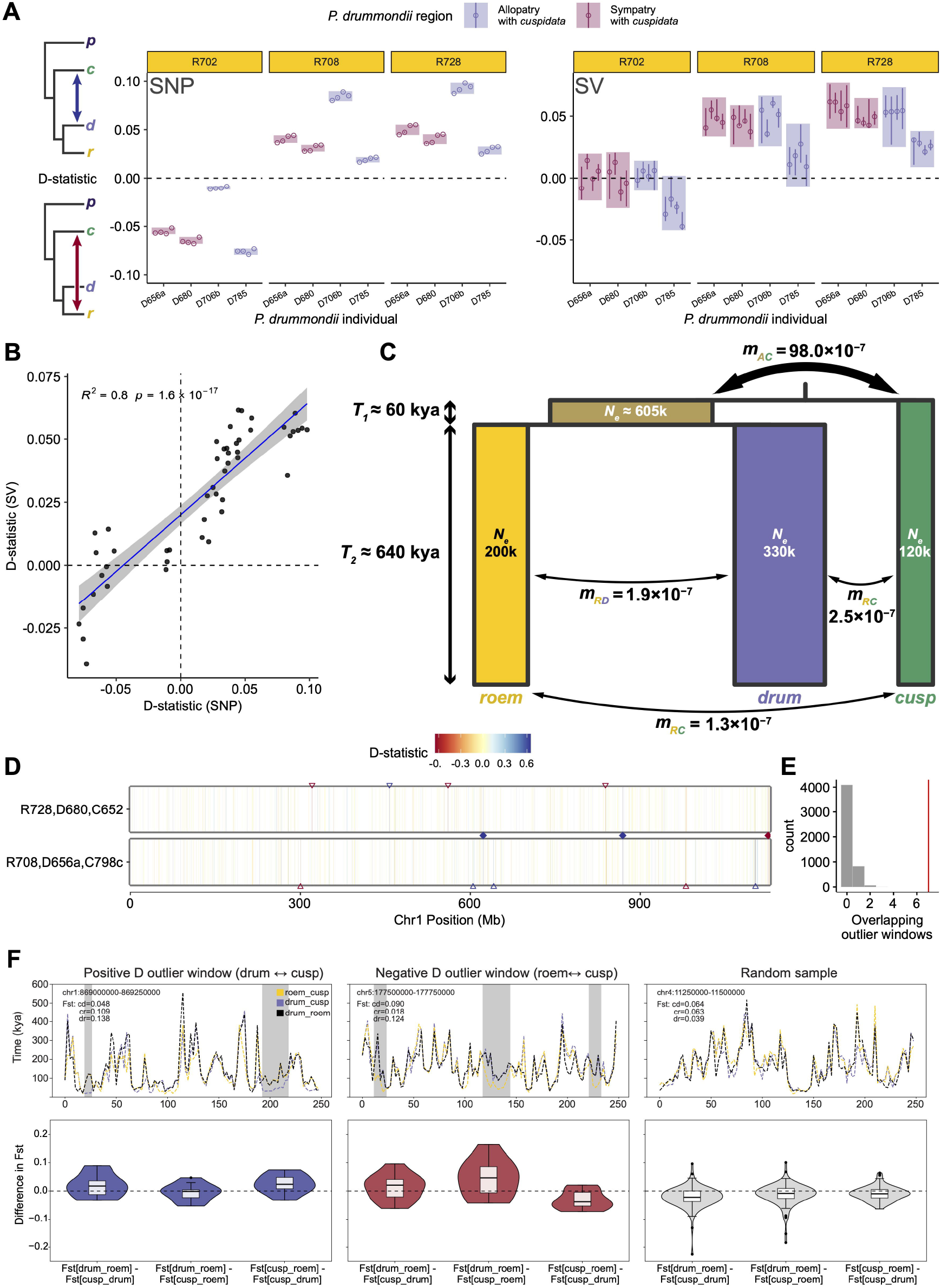
Genomic patterns of introgression across annual *Phlox*. (A) Summary D-statistic estimates from whole genome sequencing (WGS) across 48 unique three-individual trio comparisons using SNP (left panel) and structural variation (right panel). The same *P. pilosa* individual was used as outgroup in all comparisons. Positive values correspond to signals of introgression between *P. cuspidata* and *P. drummondii* while negative values indicate signals between *P. cuspidata* and *P. roemeriana*, as shown in the schematics on the top and bottom left of the graph, respectively. Comparisons are grouped by *P. drummondii* individual (x-axis) and *P. roemeriana* (yellow-bannered panels) individuals included and colored by whether the *P. drummondii* individual was sampled from regions of allopatry (purple) or sympatry (pink) with *P. cuspidata*. Points and vertical lines denote median and 95% quantile intervals, respectively, of D-statistic values across multiple replicates of allele sampling. Rectangles indicate, for a particular pair of *P. drummondii* and *P. roemeriana* individuals, minimum and maximum D-statistic values observed across all replicates and all *P. cuspidata* individuals. (B) Ordinary least squares regression of point estimates of D-statistics estimated from SNPs and structural variants. Blue line indicates best fit and grey region denotes 95% confidence intervals on the regression parameters. (C) Schematic of best-fit parameters of a three-species model of annual *Phlox* demographic history inferred by dadi using the reduced-representation sequencing (RRS) dataset. *P. roemeriana, P. drummondii*, and *P. cuspidata* lineages are depicted as yellow, purple, and green rectangles, respectively, with widths proportional to the annotated population sizes, *N*_*e*_; the ancestral lineage (beige rectangle) to *P. roemeriana* and *P. drummondii* is drawn similarly. Rectangle heights correspond to coalescence times, *T*. Symmetric, *N*_*e*_-scaled rates of migration, *m*, between all contemporaneous lineages are indicated by curved arrows with thicknesses drawn proportional to their magnitudes. (D-E) Shared local patterns of D-statistic estimates between trios {R728, D680, C652} (upper) and {R708, D656a, C798c} (lower). (D) Colored vertical lines denote D-statistic values in 250 kb windows drawn along the coordinate space (x-axis) of chromosome 1 with blue and red hues corresponding to positive and negative values, respectively. Windows with insufficient data are not depicted. Open triangles mark windows with outlier D-statistic values greater than 3 standard deviations from the mean while filled diamonds denote windows found to be outliers with the same polarity in both trios depicted. (E) Bootstrap distribution of the total number of shared outlier windows for the focal trio pair. Results from 5000 bootstrap replicates are shown; red vertical line indicates observed number of shared windows genome-wide. (F) Coalescence times and *F*_*ST*_ differences between species in outlier D-statistic windows. Upper panels show mean pairwise coalescence times between species (y-axis) from local genealogies inferred from ancestral recombination graph (ARG) reconstruction across 250 kb windows (x-axis) for representative windows showing signals of outlier positive (left) and negative (middle) D-statistic values, as well as a randomly sampled non-outlier window (right). Yellow and purple dashed lines trace pairwise coalescence times comparing *P. cuspidata* to *P. roemeriana* and *P. drummondii*, respectively. Black dashed lines denote coalescence times between *P. roemeriana* and *P. drummondii*. Grey shading highlights genomic regions in which coalescence times with *P. cuspidata* are shorter than those between sister species *P. roemeriana* and *P. drummondii* consistent with signals of introgression. Lower panels: Combination violin and box-and-whisker plots of differences in *F*_*ST*_ computed pairwise between all three species. Distributions are shown for the top ten positive (left) and top ten negative (middle) D-statistic windows as well as 100 randomly sampled non-outlier windows (right).

The above analyses were performed on nucleotide variation across *Phlox* genomes that have been masked for repetitive elements and structural variation. Given that nearly 90% of the *Phlox* genome is composed of repetitive elements, it is important to consider whether this TE-associated variation exhibits similar patterns of introgression. As described above, patterns of polymorphisms in SVs provide insights into the evolution of these genomic parasites. We therefore created a method to compute D-statistics for SVs. D-statistic values computed using SVs yield highly correlated (*p* «0.001, *R*^2^= 0.79) albeit attenuated values to those obtained from SNPs (Fig. 4A–B). When SVs are prevalent and identifiable they can be used as an alternative to SNPs to detect introgression. Yet, the discrepancy in the strength of the signal between the two forms of genetic variation motivates future research into how gene flow may impact the evolution of selfish elements across species and how evolution within species differentially impacts SVs and SNPs. Nevertheless, taken together, these results provide confidence in our emergent picture of complex admixture across the annual *Phlox* clade.

To better explore the historical patterns of divergence and gene flow among the three annual *Phlox* species, we evaluated our RRS SNP-frequency data under different migration models using the diffusion approximation method, dadi^66^. We tested three types of demographic models: (1) a “full-migration” model featuring four migration edges: pairwise between each of the contemporary lineages, as well as between the *P. cuspidata* lineage and the ancestor of *P. roemeriana* and *P. drummondii* (*r-d*-ancestor); (2) a series of “partial-migration” models in which specific migration edges were eliminated; and (3) a “no-migration” model in which only population size and divergence time parameters were retained (Table S2). In each model, migration parameters were symmetric and continuous through time.

Across all tested scenarios, the full-migration model exhibited the highest likelihood (ΔAIC = −93.74 compared to the next-best model; Fig. 4C). These results indicate pervasive gene flow throughout divergence of these species. Replicate optimizations of the full-migration scenario consistently fit better than partial- or no-migration scenarios (AIC values; Table S2). Some admixture between *P. drummondii* and the other two annual species is expected given their overlapping ranges and our observations of present-day hybrids (Fig. 1B, Fig. 2B). Notably, the full-migration model also suggests weak but persistent migration between *P. roemeriana* and *P. cuspidata* despite the species lacking any known contemporary sympatry zones. Gene flow appears to have persisted since the annual clade split from *P. pilosa* (*T* ≈ 700 kya, assuming a mutation rate of 1.0 × 10^−8^ per bp per generation) with the largest migration edge inferred between *P. cuspidata* and the *r-d-*ancestor. The smallest inferred migration edge sits between *P. drummondii* and *P. cuspidata*, which experienced reinforcement. This pattern is consistent with the evolution of flower color divergence successfully reducing gene flow between the species. Overall, our results suggest that the annual *Phlox* are a young clade with more permeable species boundaries than previously inferred based on contemporary geographic boundaries.

In addition to illuminating patterns of gene flow, the best fitting dadi models infer a large effective population size for *P. drummondii* (*N*_*e*_ = 336,564), with smaller population sizes for *P. roemeriana* (*N*_*e*_ = 200,597) and *P. cuspidata* (*N*_*e*_ = 122,412). Relative differences between the contemporary population size estimates are consistent with the species’ observed geographic ranges (Fig. 1B) and life histories: *P. cuspidata* exhibits high rates of selfing and *P. roemeriana* is an ecological specialist preferring more arid, calcareous soils^67^. We also inferred divergence between *P. cuspidata* and *P. drummondii* occurring ~1 million years ago. This value is notably smaller than previous fossil-calibrated estimates performed across *Polemoniaceae* (3.6 Mya)^68^ and *Ericales* (2.9 Mya)^69^. These differences could be due to sampling error as the fossil-calibrated estimates used limited amounts of sequence from specific genes compared to our broader genome wide sampling. Alternatively, uncertainty in the assumed mutation rate and previously observed bias in RAD-Seq technology towards lower-diversity genome regions could distort our estimates^70^. We also observed higher genetic divergence in our WGS data (Fig. S8), which is more in line with existing estimates derived from RNA sequencing^31^. We thus interpret our dating of the *P. cuspidata* and *P. drummondii* split to be a lower bound on the true value.

Given the extensive evidence of gene flow throughout the divergence of the annual *Phlox* species, we searched for patterns of variation across the genomes of these species. We performed D-statistic analyses along sliding windows of 250 kb using the WGS and reveal regions of high discordance consistent with patchy remnants of introgression (Fig. 4D). These highly discordant “outlier” windows (defined as exceeding 3 standard deviations from the mean) overlap across individuals more than expected by chance (bootstrap p-value < 2 × 10^−4^, Fig. 4E). These regions of overlap indicate certain introgressed regions of a genome may be preferentially retained across individuals post-admixture due to a combination of shared history, positive selection, and correlated recombination landscapes^71^. Across the 48 interspecific comparisons we identified 84 and 89 outlier windows exhibiting highly positive and negative D-values, respectively, in more than one comparison, and 19 outlier windows shared across more than 20 comparisons.

A pattern of consistently large D-statistic values within a particular genomic window across multiple sets of individuals can be indicative of gene flow but is also consistent with incomplete lineage sorting. To validate a history of introgression within highly shared outlier genomic regions, we ranked our outlier windows by the number of comparisons in which they were shared and used ARGweaver^72^ to reconstruct the ancestral recombination graphs (ARGs) of all our WGS samples for the ten most extensively shared positive and negative 250 kb windows (20 windows total, ranging from 20 to 48 shared comparisons, Table S3). These ARGs show variation in pair-wise divergence time between each pair of annual *Phlox* species (Fig 4F). We found that windows containing outlier signals of positive D-statistics have identifiable regions in which *P. drummondii* coalesced with *P. cuspidata* earlier than with the sister species *P. roemeriana*. Similarly, windows containing outlier signals of negative D-statistics show *P. roemeriana* coalesced with *P. cuspidata* earlier than with *P. drummondii*. These patterns of coalescence are discordant with the inferred species tree and also consistent with a history of recent gene flow but not shared ancient variation (Fig. 4F, upper panel). We quantified the enrichment of this signal by scanning the outlier windows for areas of “high” discordance, defined as contiguous regions 10 kb or larger in which the ratio of mean *drummondii*-*cuspidata* (or *roemeriana*-*cuspidata*) coalescent times to mean *drummondii*-*roemeriana* coalescent times was less than 85%. In total, we determined that 5.2% of positive and 2.8% of negative D-statistic outlier windows were composed of such regions of high discordance. As a control, we randomly sampled 100 non-outlier windows, which showed a much more modest enrichment with only 1.8% of sequences corresponding to highly discordant loci.

We corroborated our ARG inferences by estimating, for each window, *F*_*ST*_ summary statistics pairwise between each of the three annual species (Fig. 4F, lower panel). As expected, in positive outlier D-statistic windows we observed *F*_*ST*_ estimates between *P. cuspidata* and *P. drummondii* to be significantly smaller than *drummondii*-*roemeriana* values (*p* = 0.005, linear mixed modeling). In negative outlier D-statistic windows we found lower *F*_*ST*_ between *P. roemeriana* and *P. cuspidata* relative to the other possible comparisons. Non-outlier windows showed *F*_*ST*_ patterns more consistent with no or little admixture such that *P. cuspidata* exhibited similar levels of *F*_*ST*_ with both *P. roemeriana* and *P. drummondii*, on average.

The presence of a dark red corolla variant of *P. drummondii* in central Texas has been appreciated since at least the 1940s^24,25^, and its role in limiting hybridization with *P. cuspidata* through sympatric reinforcement has been studied since the 1980s^27,73^. More recently, sequence analyses have added genetic support for past introgression between this focal species pair^31,32^. However, our results paint a more sophisticated story—namely high gene flow between the rapidly speciating annuals in a highly intermixed ancestral population followed by continued incidents of gene flow between not just two but all three lineages. These results are robust with multiple introgression signals from both reduced-representation and whole genome long-read sequencing, and from both single nucleotide polymorphisms and structural variants. Yet our findings also highlight the complex genealogical relationships of the system and the need for similarly complex models to account for them. For instance, our demographic modeling ignores the derived selfing behavior of *P. cuspidata*, which imposes specific expectations for asymmetry in introgression^74–76^. Our genome assemblies allow us to, for the first time, identify specific genomic regions derived from recent admixture events, prompting new questions about the evolutionary forces—selection, drift, and recombination—that have preferentially maintained these sequences.

## Conclusion

Our high-quality chromosome level *Phlox* genome assemblies, along with dense datasets of genetic variation, deepen our understanding of the complex dynamics underlying plant evolution and the speciation process. Decades of work on patterns of diversification in *Phlox* has rendered this complex of Texas annuals instrumental to our understanding of how species form and maintain their boundaries; yet previous lack of genomic resources for this system, and other large, complex plant genomes, has limited our advancements. The three annual *Phlox* species, *P. drummondii, P. roemeriana*, and *P. cuspidata*, have a tangled evolutionary history involving extensive gene flow throughout divergence between all three species. Evidence of this gene flow is heterogenous across geographic space and across chromosomes suggesting other forces of selection and demography are shaping patterns of genetic variation as well. Nearly 90% of a *Phlox* genome is composed of repetitive elements leading to abundant structural variation and complex genome structure. We untangle this complexity by associating TEs with SVs and revealing clear phylogenetic signal in this abundant variation. The variation in SVs also harbors consistent evidence of gene flow between *Phlox* species indicating this often-masked variation can be informative for evolutionary inference. In these *Phlox* species, striking patterns of phenotypic divergence (such as dark red flower color) are attributed to forces of selection and gene flow during speciation, and yet we find the dominant patterns of genomic variation in the annual *Phlox* species are shaped by life-history strategy, mating system, and abiotic factors. By assembling some of the largest chromosomes for any species, and unmasking repetitive and structural variation, we provide a roadmap for evolutionary inference of complex whole genome data with this non-model system.

## Methods

### Plant collection and propagation

All *Phlox* plants in this study are derived from seeds collected from wild plants over a period of 3 years from the southern United States (primarily Texas, see Table S1 and S4 for collection details). The plants were grown under controlled greenhouse conditions with 16 hours of supplemented daylight and temperature between 18 °C and 25 °C at the Arnold Arboretum of Harvard University. Seeds were induced to germinate by soaking in 500 ppm gibberellic acid in water for 48 hours and subsequently cold stratifying planted seeds at 4 °C for ten days. Plants were grown in 4-inch pots with Pro-Mix HP Mycorrhizae potting media (Quakertown, PA, USA). Young leaf and bud tissue was collecting and stored at −80 °C for DNA extraction. Tissue from leaf, stem, pistil, pollen, and anthers were collected and immediately flash frozen in liquid nitrogen then stored at −80 °C for RNA extraction.

### Long-read sequencing

To stimulate meristematic tissue growth prior to nuclear extraction, plants were placed in dark chambers for 48 hours. Approximately 1.5 g of fresh meristem was collected from each plant, flash-frozen in liquid nitrogen, and ground into a dry frozen powder using mortar and pestle. Powderized tissue samples were suspended in 25 mL HB buffer (50 mL 1X HB buffer + 150 µL β-mercaptoethanol at concentration). The resulting slurry was filtered through 22–25 µm Miracloth™, and the filtrate was centrifuged at 2,400 g for 20 minutes at 4°C to separate the nuclei. The pellet was resuspended to 25 mL of HB buffer using a wide bore pipette tip. Filtration and resuspension of nuclei was repeated 2–3 times for each sample before a final round of resuspension in 1 mL of HB buffer and apportioned into five 200 µL aliquots per sample, which were frozen at −80°C.

We extracted high molecular weight (HMW) DNA from isolated nuclei. Each aliquot of frozen nuclei was resuspended in 750 µL of CTAB buffer and warmed in a 55°C water bath with occasional inversion. Next, 950 µL of 24:1 chloroform:isoamyl-alcohol solution was added to the nuclei solution, and the sample tubes were manually agitated to achieve mixing. The mixture was separated with 15 minutes of centrifuging at 5,000 g, and the aqueous phase was retained. Samples were incubated at 37°C for 30 minutes with 10 µL of RNAse A before a second chloroform wash-and-centrifuge step was performed.

To precipitate DNA, 54% volume of isopropanol (~1,350 µL) and 10% volume of 4°C Sodium Acetate (3M, pH 5.2, ~250 µL) were added to each sample. Samples were allowed to sit undisturbed for 15–60 minutes to allow precipitation to occur prior to centrifuging the samples at maximum speed for 25 minutes to pellet the precipitate. The supernatant was removed, and the pellet was washed with 1 mL of 4°C 70% ethanol. An additional two rounds of centrifugation for 10 minutes and 1 mL of 4°C 95% ethanol wash of the pellets were subsequently applied to clean the pellet. The pellets were centrifuged a final time for 10 minutes and allowed to airdry for 10–15 minutes following removal of the supernatant. Samples were resuspended in 100 µL of TE solution pH 8 and stored at 4°C for multiple days to ensure DNA product had fully dissolved into solution. Standard Nanodrop, Qubit, and gel electrophoresis, procedures were used to quantify and quality control the finalized HMW DNA samples.

Samples were sequenced on the Oxford Nanopore Technologies PromethION at the Bauer Core Facility at Harvard University. For population resequencing samples, we took advantage of newer “super” high-accuracy basecalling models and re-basecalled our data using Guppy v6.4.6. We used the configuration file ‘dna_r9.4.1_450bps_sup_plant.cfg’ which is a super-accuracy model optimized for plant sequences. Three of the samples (702-2-3-1, 706b-21-c, and Hopkins_1704_09) were not re-called, as they were sequenced using a newer sequencing chemistry and basecalled using the default high accuracy model in MinKnow.

### Short-read sequencing

We sequenced the genome of reference *P. drummondii* individual MPI409 using short-read sequencing technology to supplement our long-read data. To this end, we extracted DNA from bulk meristem and leaf tissue and prepared libraries using the Nextera Illumina Library Kit (indexes i5=S513 and i7=N718). The libraries were size-selected to a target fragment size of 300-500 bp and subjected to 150 bp paired-end sequencing on the Illumina NovaSeq 6000 platform generating 296 Gb of total sequence.

We additionally employed reduced-representation sequencing (RRS) to sample genome-wide variation across 892 *Phlox* samples collected as seed from wild populations, and 228 individuals from a genetic mapping family used to generate a genome-wide linkage map. DNA was extracted from bud meristem tissue from the plants of interest. We prepared genomic libraries using a previously described double digest restriction-site-associated DNA (ddRAD) sequencing protocol^28^, which subsequently underwent 150 bp paired-end sequencing on the Illumina NovaSeq 6000 platform.

Raw ddRAD sequence reads were demultiplexed using the Stacks software suite v2.65^77,78^. First, we processed the forward reads using a bespoke script to clip the unique molecular barcode from each sequence of the raw FASTQ and move it to the header. We then proceeded to demultiplex each FASTQ file pair using process_radtags with the --paired flag set and passing in the relevant sample barcodes using the -b flag. Duplicates were removed from the demultiplexed FASTQs—one pair for each sample—using the clone_filter tool with the --null_index flag set and --oligo_len_1 set to the length of the unique molecular barcode. As a final processing step, we applied the Trimmomatic tool v0.39^79^ to remove poor quality reads or bases and stray Illumina adapter sequences.

### *P. drummondii* genome assembly

Using the ONT long-reads and Illumina short-reads obtained from individual MPI409, we *de novo* assembled a *P. drummondii* genome. We followed a typical process of assembly with the Flye algorithm followed by multiple rounds of polishing and scaffolding to obtain a final haploid assembly^80^.

Initial assembly was done using Flye v. 2.8.3^81^ with a total of 241 Gb—approximately 35X coverage, assuming a 6 Gb haploid genome^35^. We performed the assembly on a dedicated node with 1 TB RAM using 30 threads for a total assembly time of 78 days. To remove assembly artifacts and duplicate contigs, we ran the Purge Haplotigs^82^ pipeline on the draft assembly; we elected to retain contigs marked by the pipeline as “repeats” as removing them resulted in a significant loss of BUSCO genes. To correct errors, we subsequently performed a “self-polish” of the assembly using the Nanopore long-reads and the GPU version of Medaka v1.2.6^83^. Reads were aligned to the reference using Minimap2, and medaka polish was run in parallel batches to improve run time.

To increase contiguity the draft assembly was optically mapped with Bionano at the Genome Technology Access Center at the McDonnell Genome Institute at Washington University, St. Louis, USA. Libraries were prepared from flash-frozen tissue and run on two Bionano Saphyr flow cells. Scaffolding was performed on the polished draft assembly using the default Bionano hybrid scaffolding workflow while allowing for sequence cuts in the conflict resolution algorithm. Preliminary comparison of the Bionano-scaffolded assembly to linkage information (see next paragraph) indicated a single large misassembly on “Super-Scaffold_33623,” which we resolved by splitting the scaffold at positions 109,426,292 and 109,426,293. This procedure produced an improved assembly having a scaffold N50 > 6 Mb.

To further increase contiguity, we performed a second round of scaffolding incorporating a linkage map. The linkage mapping population consisted of two P0 parental individuals, a single F1 offspring, and 225 F2 siblings, which were produced from an autogamous cross of the F1 individual. Our reference genome individual was an individual of this mapping population. We performed RRS on the 228 total individuals (see “Short-read sequencing”) and the pre-processed reads were aligned to the Bionano assembly sequences using BWA-MEM v0.7.17 (parameters -k 15 -r 1.3 -T 20). SNPs were called using the *Stacks* suite (see “Calling SNPs from reduced-representation sequences”), and the map was constructed using the Lep-Map3 suite v0.5^84^. Since Lep-Map3 was designed for allogamous settings, we configured the pedigree file to account for the selfed F1 by providing a “dummy” male and “dummy” female individual whose parents were specified as the P0 individuals. These two dummy individuals were assigned as parents to the 225 F2 individuals, and the actual F1 individual was removed from the analysis. We passed the modified pedigree of the mapping population and the VCF file of SNPs to ParentCall2, a Lep-Map3 module that generates parental genotype markers. Markers were filtered using Filtering2 module (parameters removeNonInformative=1, dataTolerance=0.0001, MAFLimit=6, missingLimit=0.5) and subsequently assigned to linkage groups using SeparateChromosomes2 (parameters lodLimit=50, theta=0.0013, sizeLimit=50, and grandparentPhase=1). We used JoinSingles2All to assign unassigned markers to the main linkage groups using the same lodLimit and theta settings as in the previous step. Finally, we applied the OrderMarkers2 module to output phased markers ordered within each linkage group (parameter outputPhasedData=1). Our mapping pipeline produced seven large linkage groups, corresponding to the seven chromosomes. Markers from the seven linkage groups were reformatted and combined to produce a final genetic map in BED format, as required by ALLMAPS, which was used to scaffold the final assembly (Tang et al. 2015).

The final scaffolded assembly was highly contiguous, with a scaffold N50 of 406Mb. To correct any remaining errors, we aligned Illumina short-read data against the reference using Bowtie2^85^ and polished the assembly using the Illumina short reads and HyPo v1.0.3^86^. The final BUSCO completeness score using the viridiplantae_odb10 database was >99%.

### Assembly of other *Phlox* species

For the three remaining *Phlox* species (*P. cuspidata, P. roemeriana, P. pilosa*), we used Oxford Nanopore Technologies PromethION to sequence 91–225 Gb from each individual. Base-called ONT sequences were initially assembled using Shasta v0.11.1 with 1600GB of RAM and 32 threads on a single dedicated node, and with the --memoryMode anonymous --memoryBacking 4K settings. We used the ‘Nanopore-Plants-Apr2021.conf’ configuration preset, which is intended to improve assembly for large, complex plant genomes. Total assembly time was between 2 and 4 days per species.

Initial contig assemblies were then scaffolded against our highly contiguous *P. drummondii* reference using Ragtag v2.1.0^87^ with the following parameters for the Minimap2 alignment step: --mm2-params='-x asm10 -I 20G'. The scaffolded assemblies were polished using self-correction with Medaka v1.7.2 (GPU version)^83^. Reads were aligned to scaffolds using Minimap2 and polished using medaka polish in parallel batches to improve runtime.

### Multiple sequence alignment

We aligned the scaffolded genomes for *P. drummondii, P. cuspidata* and *P. roemeriana* against the *P. pilosa* reference using wfmash v0.10.5-4-g49f9b10^88^. Due to their large sizes and repetitive makeup, we first did an approximate alignment using the following wfmash parameters: -m -p 90 -t 8 -c 100k. We then split the approximate alignment into ten batches using the split_approx_mappings_in_chunks.py script from wfmash. We refined each alignment chunk in parallel using the -i option in wfmash, and combined chunks to make a final alignment.

To generate whole genome alignments between our finished reference genomes, we used AnchorWave^10,36^, as we found it best suited to large, repeat-dense plant genomes, though it is only capable of pair-wise alignment. AnchorWave uses conserved CDS regions as “anchors” for initial alignment to create locally collinear blocks. We extracted the CDS annotations from our *P. drummondii* gene annotation and mapped the CDS to the *P. drummondii* genome as well as each of our other species using Minimap2 with the ‘splice’ option to generate CDS anchors for alignment. Using *P. drummondii* as our reference, we then ran anchorwave proali on each genome to generate pairwise alignments with *P. drummondii*. At time of analysis, the multiple alignment format (MAF) output of AnchorWave contained an error in the representation of start positions for negative strand alignments, reporting coordinates for the positive strand instead^89^; we applied a bespoke awk script to the MAFs to correct this issue.

### Repeat identification and annotation

We generated a library of repetitive elements by identifying them *de novo* from the reference *Phlox drummondii* genome using RepeatModeler^90^, which includes several repetitive element-finding programs including RECON, RepeatScout, LtrHarvest and Tandem Repeat Finder. A Singularity image of RepeatModeler is provided by the Dfam Transposable Element Tools consortium^91^. First, we constructed a database of the reference genome using the BuildDatabase function, then ran RepeatModeler with the -LTRStruct option for the LTR discovery pipeline.

We also ran the Satellite Repeat Finder (SRF) pipeline to better identify tandem repeats in the genome^92^. First we identified kmers in the genome using KMC^93^ with the parameters -m60 -k151 -t16 -ci25 -cs100000 and used the kmc_dump function to generate a list of counts for each kmer. We then used the counts with SRF to generate a FASTA file of tandem repeats.

### Gene annotation

We annotated the *P. drummondii* genome using two bulk RNA-Seq datasets collected from multiple tissues including leaf, stem, pistil, pollen, and anthers. We performed the transcriptome alignment, assembly, and annotation steps using a Snakemake workflow developed by the Harvard Informatics group^94,95^. The RNA-Seq reads were first trimmed using trim-galore^96^ with parameters --paired --retain_unpaired --phred33 --length 36 -q 5 --stringency 1 -e 0.1. The trimmed reads were aligned to the *P. drummondii* reference genome with HiSat2 v2.2.1^97^, and the resulting alignments were sorted using samtools sort v1.13^98^. The sorted alignments were individually passed as inputs into StringTie v2.1.2^99^ for transcriptome assembly and then merged using the stringtie --merge module. We finally applied the Transdecoder^100^ pipeline v5.5.0 to process the assembled transcriptome GTF and predict coding DNA sequence regions throughout the genome; a custom Python script was used to integrate non-coding RNA sequences into the final annotation.

### Calling SNPs from long-read sequences

In addition to our *P. drummondii* reference genome individual, we used ONT to long-read sequence a total of 12 annual *Phlox* individuals. We performed SNP calling on our 12 long-read sequenced individuals (Table S1) against both the *P. drummondii* assembly as well as the *P. pilosa* assembly and used the resulting datasets for different analyses (e.g., ingroup reference for gene annotation comparisons, outgroup reference for population genetics, D-statistics). The ONT FASTQ sequences were aligned to a given reference assembly using Minimap2 v2.26-r1175^101^ in ONT mode with the parameters -ax map-ont --secondary=no -I 30g -K 1G -L. (Note that both the *P. drummondii* and the *P. pilosa* reference assemblies require more RAM than the average genome to index, hence the -I 30g argument to prevent multiple mappings against a multi-part index.) We used *samtools sort* to sort the reads in the resulting BAM files and samtools merge to combine the files associated with each sample, resulting in a single sorted BAM file for each of our 12 samples.

ONT long-read sequencing data exhibits a higher per-base-per error rate than short-reads. To account for this error in our downstream analyses we opted to use a genotype sampling scheme. For every individual we calculated genotype probabilities at all covered sites using bcftools mpileup (v1.21) in ONT mode with the arguments -X ont -a AD,DP. With a custom Python script, we then used these probabilities to sample and write genotypes to a VCF. If the sampled genotype included the missing/deletion allele “*”, then the genotype was set to missing. In total we performed five sampling replicates per individual and data was merged using bcftools merge.

### Calling SNPs from reduced-representation sequences

We mapped our 892 preprocessed RRS samples to the *P. pilosa* assembly using the BWA-MEM algorithm v0.7.17^102^. With Stacks suite v2.65^77,78^ we identified RAD loci using gstacks (parameters --rm-unpaired-reads) and called variants at all sites using the populations module in -- vcf-all mode. The “population” groupings required by *Stacks* were defined using species assignment.

To consider the impact of filtering criteria on our results, we constructed a Snakemake pipeline to apply different filtering thresholds during the *populations* calling step. Specifically, we required RAD loci to be present at thresholds of at least 10%, 50%, or 70% of individuals in the population and overall (i.e., arguments --min-samples-per-pop and --min-samples-overall). We also applied different tiers of missing data thresholds using PLINK v1.9^103,104^, filtering out individuals with greater than 25%, 50%, 75% or 90% missing SNP data.

### Genetic variation along the genome

We computed population genetic statistics *π* and *F*_*ST*_ in windows along the *P. drummondii* genome using our five replicate whole genome SNP call-sets (see “Calling SNPs from long-read sequences”). Statistics were computed on each replicate using the tool pixy v1.2.10.beta2 in 1 Mb windows^105,106^. For each window, we plotted the median value in a circular figure using the circlize package in R^107^. The circlize package was used to plot gene annotation data in 5 Mb windows in a similar manner.

### Calling structural variants

To call structural variants, we adopted a consensus approach where long-reads from twelve individuals across the three annual *Phlox* species were mapped to the *P. pilosa* reference and evaluated using three different SV calling programs. We organized the pipeline into a Snakemake workflow, summarized as follows: five long-read samples from *P. drummondii*, four from *P. cuspidata*, and three from *P. roemeriana* were mapped against the *P. pilosa* reference using Minimap2, as *P. pilosa* is the outgroup to the three other species (see “Calling SNPs from long-read sequences” above). We then used the BAM files as input to three different SV calling programs: SVIM^108^, cuteSV^109^ and Sniffles2^110^. Calls for each sample were then merged using SURVIVOR^111^, with the following command: SURVIVOR merge {params.text} 1000 3 1 1 0 50 {output}, which merges SVs within 1000bp of each other that are identified by all three callers (i.e. the intersection of the three sets of SVs). We then filtered the SVs to remove generic breakpoints (BNDs) and translocations, as well as any SV > 100kb in size.

We merged SVs calls across all twelve samples using SURVIVOR merge, this time retaining an SV if it was present in any one of the samples: SURVIVOR merge {params.text} 1000 1 1 1 0 50 {output}. Finally, to genotype each SV for each sample, we provided the combined set of SVs to Sniffles2 using the --genotype-vcf option for each sample, which we then combined into one final set of SVs. Note that while the reference individual *P. drummondii* was included in SURVIVOR merging, we ultimately omitted this individual from downstream analyses as the sequence coverage was much higher

### Tallying transposable element and structural variant associations

To better understand TE dynamics, we associated different TE families with SVs. We prepared a FASTA file containing sequences of identified SV alleles (see “Calling structural variants”) and passed this file through our RepeatModeler pipeline (see “Repeat identification and annotation”), resulting in a summary table of identified SVs and the TE families detected within their sequence tracts. For some SVs, the SV caller did not provide clear sequence information, and those SVs were therefore not included in the dataset. SVs with missing genotypes across the eleven samples were also removed. In total, 267,735 SVs were retained. For each TE family, we tabulated a “TE-SV count” *N*_*Jam*_, for the number of occurrences of that family found among a particular set of SVs, *N*_*Jam*_ [sv]. We emphasize that TE-SV counts are tallied conditional on a particular set of SVs and notate as such. We computed TE-SV counts for all the SVs observed in each of the annual species (*N*_*Jam*_ [sv_*drum*_], *N*_*Jam*_[sv_*roem*_], and *N*_*Jam*_ [sv_*cusp*_]) as well as for SVs observed *uniquely* in each species (*N*_*Jam*_ [sv_*drum,unique*_], *N*_*Jam*_ [sv_*roem,unique*_], and *N*_*Jam*_ [sv_*cusp,unique*_]). We additionally counted the number of copies of each TE family in the repeat annotations of each species’ reference genome (*N*_*Jam*_ [*Ref*_*drum*_], (*N*_*Jam*_ [*Ref*_*roem*_], and (*N*_*Jam*_ [*Ref*_*cusp*_]).

To estimate correlations between TE-SV counts, we used generalized linear modeling (GLM). Using the glm.nb function from the MASS package in R^112^, we performed negative binomial regression with an identity link function using OLS estimates (floored at 0) for the regression parameters as starting values in the linear predictor. Pseudo-*R*^2^values were computed as 1 minus the ratio of model deviance to the null deviance.

We observed the correlation between *N*_*Jam*_ [*sv*_*cusp,unique*_] and *N*_*Jam*_ [*sv*_*drum,unique*_] to be weaker than that of *N*_*Jam*_ [*sv*_*cusp*_ ] and *N*_*Jam*_ [*sv*_*drum*_], based on pseudo-*R*^2^values. As *sv*_*sp,unique*_ is a subset of *sv*_*sp*_, we used a bootstrap resampling procedure to estimate the likelihood of observing this reduction in correlation by chance. For a given species we subsampled the SVs present (*sv*_*sp*_) without replacement requiring the size of each subsample to be equal to the number of unique SVs (*sv*_*sp,unique*_). We performed 500 rounds of subsampling in each of *P. drummondii* or *P. cuspidata* and estimated pseudo-*R*^2^values using GLM as described above, resulting in a distribution of pseudo-*R*^2^values.

### Phylogenetic tree inference

We inferred phylogenetic trees from our twelve long-read WGS samples using RAxML-NG-MPI^113^. To ensure that input sequences were putatively neutral and that reduce the likelihood of erroneous SNP calls, we restricted input sequences to genomic regions that were annotated as non-coding (including 1 kb of flanking sequence) and non-repetitive. We also required at least 10 or more reads (DP format field) and presence in all twelve samples for a site to be considered. We used bcftools to filter the raw VCFs (see “Calling SNPs from long-read sequences” above) and vcf2phylip^114^ to convert the result into PHYLIP format. We ran raxml in multi-threaded mode with the discrete Gamma model with four categories (--model GTR+G), 100 bootstrap trees (--bs-trees 100), and 10 parsimony and 10 random starting trees (--tree pars{10},rand{10}); we rooted the trees using the *P. pilosa* sample as the outgroup (--outgroup ppil_Hopkins_1704). Tree figures were drawn in R using ggtree^115^.

### Population genetic and ancestry inference

We processed our RRS SNP calls (see “Calling SNPs from reduced-representation sequences”) using PLINK v1.9^103,104^ in preparation for principal component analysis (PCA). For PCA, we performed LD pruning using the --indep-pairwise function in PLINK with arguments “50 10 0.1”— i.e., removing variants with pairwise *R*^2^values greater than 0.1 in sliding windows of 50 variants with a step size of 10 variants. We additionally removed variants with a minor allele frequency less than 2.5% (--maf 0.025) and exceeding a Hardy-Weinberg equilibrium test by a p-value of 10^−16^ (--hwe 1e-16). To mitigate the effects of missing data on our PCA we used a missing forest scheme implemented in the missForest package in R^116^ to impute uncalled genotypes. We used the frequencies of the three genotypes in each sample as priors, allowed 100 trees per sample, and took the floored square root of the number of samples as the value of the parameter mtry. We allowed for a maximum of 10 iterations in imputation. PCA was performed using the –pca function in PLINK and results were plotted in R using the graphics package ggplot2^117^.

We additionally performed an analysis of molecular variance (AMOVA)^118^ on the variant set used as input for our PCA. A separate AMOVA was applied to each species with population as the grouping variable. AMOVAs were performed using the function poppr.amova from the poppr package v2.9.6 in R^119^.

We prepared the RRS SNP calls for ancestry estimation in a similar manner as for PCA, pruning SNPs for LD (--indep-pairwise 50 10 0.1) and for Hardy-Weinberg disequilibrium (--hwe 1e-16). We inferred ancestry components using ADMIXTURE v1.3.0^43^ with group numbers (parameter *k*) of 1 through 8, inclusive (Fig. S9). For each *k*-value we performed three reseeded replicates and retained the one with the highest log-likelihood.

To infer contemporary migration patterns we applied the Estimation of Effective Migration Surfaces (EEMS) package to our RRS SNP data^48,120^ using the same PLINK filter settings as in our admixture analysis. We used the bed2diffs and coord_from_fam.sh tools to convert our PLINK dataset into formats acceptable by EEMS. We set the number of demes to 350 and allowed for 10,000,000 MCMC iterations with 1,000,000 burn-in iterations and a thinning interval of 9,999. The outer boundaries of the habitat were selected *ad hoc* so as to contain all populations across the three annual species, and the EEMS algorithm was executed using the runeems_snps tool. Resulting surfaces were plotted using the reemsplot2 library^121^.

To assess isolation by distance we computed pairwise Weir-Cockerham *F*_*ST*_ estimates between populations using the corresponding function from Python package scikit-allel^122,123^. Populations comprised of a single individual were ignored. We regressed our pairwise *F*_*ST*_ estimates against pairwise population Haversine distance using ordinary least squares (OLS). For *P. drummondii*, we additionally examined isolation by distance in only the subset of population pairs that involved exactly one population in sympatry with *P. cuspidata* and one population in allopatry. To understand if the regression slope of comparisons of allopatric versus sympatric *P. drummondii* populations deviated significantly from null expectation we compared the observed slope against slopes obtained from 500 bootstrap resamplings of the *P. drummondii* pairwise *F*_*ST*_ dataset.

### Demographic inference with dadi

To obtain model-based estimates on annual *Phlox* demographic history and introgression over time, we relied on dadi^66^. For dadi inference we used RRS data from 722 *P. pilosa-aligned* samples: 161 cuspidata, 133 roemeriana, 428 drummondii. After a grid search, we projected the SNP data down to 32, 26, and 100 individuals, respectively, to maximize the number of segregating sites. We sampled one SNP per RAD locus. We recovered 7767 segregating sites in a folded spectrum and designated extrapolation grid points of 140, 180, and 220. We tested three different three-population models from the dadi_pipeline toolkit from^124^: no_mig, split_symmig_adjacent, split_symmig_all. For each model, we performed three separate optimizations consisting of four folds and 100 total replicates. Each optimization routine consisted of: round 1, 3-fold parameter perturbation, 10 replicates; round 2, 2-fold parameter perturbation, 20 replicates; round 3, 2-fold parameter perturbation, 30 replicates; round 4, 1-fold parameter perturbation, 40 replicates.

### ABBA-BABA testing and ancestral recombination graph inference

Using our *P. pilosa*-aligned WGS SNP data, we computed f-branch values as in Malinsky et al.^65^. We assigned the single *P. pilosa* individual as the outgroup and treated each sampled individual as a unique leaf, under the phylogenetic relationships inferred using RAxML (see “Phylogenetic tree inference” above). We filtered SNPs for complete genotyping data (AN == 12) and assessed a minimum coverage of at least 5 reads (DP > 5) to retain a genotype. In practice, we calculated f-branch values using the Dsuite toolset^125^ and plotted the results using the included plotting script dtools.py.

To further explore allele sharing patterns, we calculated D-statistics on our *P. pilosa*-aligned WGS SNP and SV data using a quartet approach to maximize data availability. In each comparison a single representative individual of each species was chosen. At each variant site we sampled a single allele from each of the four individuals’ genotypes and categorized them by topological patterning: ABBA, BABA, or neither. D-statistic values were then computed using the total ABBA and BABA pattern counts^126^. The above sampling-and-counting procedure was repeated five times for each quartet comparison in both SNPs and SVs. For SNPs, each replicate was performed on a different genotype sample set (see “Calling SNPs from long-read sequences”), and we also required a minimum coverage of at least 5 reads (DP > 5). We computed D-statistics for each quartet possible (48 total) and regressed median SNP and SV D-values using OLS.

To compute windowed D-statistics we divided the *P. pilosa* genome into 250 kb non-overlapping windows and performed the same allele sampling procedure as above on our SNP genotypes. We pruned the windows, eliminated those with fewer than 1000 informative sites available (i.e., sites with an ABBA, BABA, or BBAA pattern) or with 50% or more genotypes missing across all sites. Using the median across replicates as point estimates for the D-values for each window, we identified “outlier” windows that showed an excessively high or low D-value greater than three standard deviations from the mean. For trios {R728, D680, C652} and {R708, D656a, C798c} we tested the likelihood of the observed number of overlapping outlier windows using a bootstrap resampling approach; windows were required to have the same sign be considered coincident.

We ranked our collection of outlier windows by the number of times a particular window appeared across all 48 quartet comparisons. The top ten windows most frequently found to be positive or negative outliers (Table S3) were earmarked for downstream ancestral recombination graph (ARG) inference. We also sampled 100 random non-outlier windows as a control for ARG inference.

For each window, the corresponding VCF file was extracted and used as input for ancestral recombination graph (ARG) inference using ARGweaver^72^. Windows were processed individually to ensure tractability of ARG inference and to allow independent replication across genomic regions. For each window, we used the function arg-sample with parameters mutrate=1e-8, compress-seq=10, maxtime=1e6, mask-cluster=2,5, and iters=5000. Samples were recorded every 10 iterations. For analysis, we examined the .smc file produced in the last iteration for each window, as recommended by Guo et al.^127^.

To summarize local genealogical variation across each 250 kb window, we sampled 100 evenly spaced locations. At each location, we extracted the local genealogy with the Python package toytree^128^, and using toytree functions we calculated mean coalescence times for individuals between each pair of species (*drummondii*-*cuspidata, roemeriana*-*cuspidata, drummondii*-*roemeriana*).

For each window, Weir-Cockerham *F*_*ST*_ estimates between species were calculated using a Python script as described above (see “Population genetic and ancestry inference”). We tested whether *F*_*ST*_ between *P. drummondii* and *P. roemeriana* was systematically greater than *F*_*ST*_ between non-sister lineages as would be expected in a recent admixture scenario (i.e., *cuspidata*-*drummondii* for positive outlier windows and *cuspidata-roemeriana* for negative outlier windows). We used a linear mixed model:

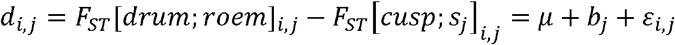

where *i* denotes the window index, indicator *j* ∈ 0,1 denotes the type of outlier window (positive or negative), and *s*_*j*_ denotes the species (*drummondii* if positive, *roemeriana* if negative). Variables *b*_*j*_ and *ε*_*i,j*_ capture random change in the mean difference for window type *j* and residual error, respectively. We tested the alternative hypothesis of mean *µ*> 0.

## Acknowledgements

We thank Christina Steinecke, Austin Garner, Anna Feller and the rest of the Hopkins lab for suggestions and comments throughout this project. Plant care was provided by the Arnold Arboretum of Harvard University Greenhouse Staff. Thank you to Faculty of Arts & Sciences Research Computing and the Bauer Core Facility at Harvard University for supporting the computational aspects of this work. This study was funded by National Science Foundation Award DEB-1844906 and National Institutes of Health Award 1R35GM142742 to RH.

## Data availability

The assembled *Phlox* genomes and associated sequencing data are available on NCBI under BioProjects PRJNA1219593 and PRJNA1223900. WGS and RRS data are available at BioProjects PRJNA1444519 and PRJNA1449435, respectively. BioProject PRJNA1450658 contains RRS data from the linkage mapping population. Scripts for analysis and reproduction of the figures may be found at: https://github.com/HopkinsLab/phlox_genome_scripts.

