## Supplementary Figures for "Unmasking the large and highly repetitive genome of *Phlox* reveals a complex evolutionary history of speciation with gene flow"

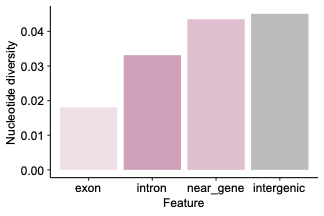

**Figure S1. Nucleotide diversity (**$\boldsymbol{\pi}$**) is reduced in coding regions.** Nucleotide diversity estimates are shown for annotated exon, intron, near-gene, and intergenic regions of the *P. drummondii* genome. Estimates were drawn from four whole genome sequenced individuals. Near-gene elements were defined as the 1kb region up- or downstream of a gene feature.

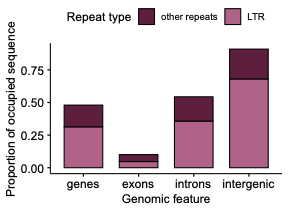

**Figure S2. Repeat occupancy across genomic features.** Bar plots of proportion of different annotated features of the genome consisting of repeat/TE sequence. Bars are stacked to show relative proportions of long terminal repeats (LTR, light pink) compared to other repeat types (dark pink).

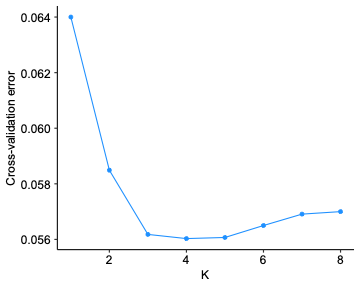

**Figure S3. Cross-validation results for the number of populations,** $\boldsymbol{K}$**, in ADMIXTURE.** Cross-validation results for parameter $K$ (x-axis), the number of proposed ancestral populations to fit in the ADMIXTURE algorithm. We tested values of $K$ ranging from 1 to 8. While the parameter is minimized at $K=4$, the majority of improvement in cross-validation error was achieved at $K=3$.

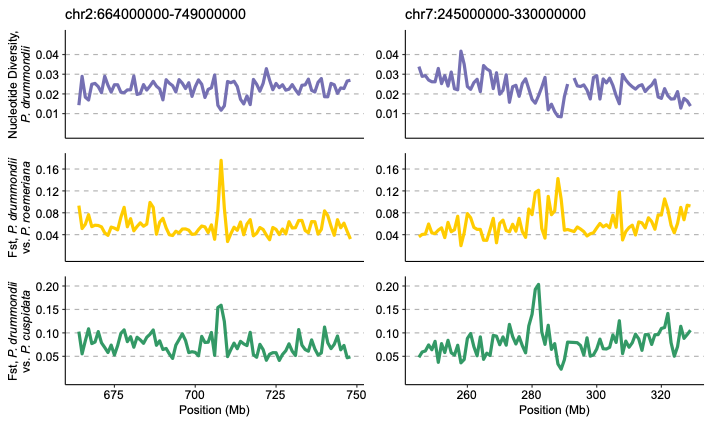

**Figure S4. Regions of exceptional divergence.** Focused view of Fig. 1 on 85 Mb-wide regions of chromosome 2 (left panel) and chromosome 3 (right panel) of the *P. drummondii* genome harboring notable patterns of divergence across the annual *Phlox*. Purple lines denote nucleotide diversity ($\pi$) in *P. drummondii* while yellow and green lines denote $F_{ST}$ between *P. drummondii* and *P. roemeriana* and *P. cuspidata*, respectively. All statistics are estimated in 1 Mb windows.

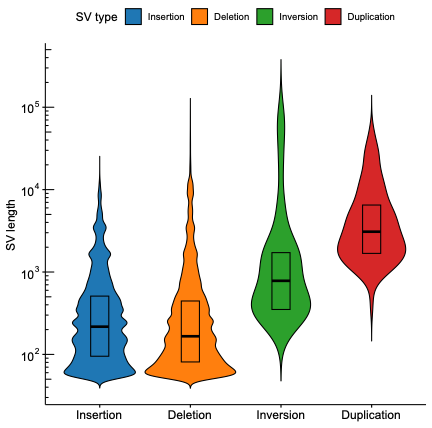

**Figure S5. Structural variant (SV) size distributions.** Violin plots of SV length (y-axis, log-scaled) distributions of insertions, deletions, inversions, and duplications. Boxes delimit interquartile ranges for each distribution, and thick horizontal lines denote medians.

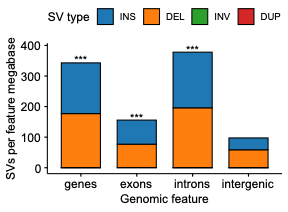

**Figure S6. Structural variant (SV) occupancy across genomic features.** Stacked bar plots SV counts in different annotated features of the genome normalized by the total sequence size of each feature in Mb. Insertions (INS, blue) and deletions (DEL, orange) predominate, and thus, inversions (INV) and duplications (DUP) are not visibly rendered. Asterisks (***) indicate feature categories for which SVs are significantly overrepresented relative to the amount of sequence available ($p<0.001$, binomial test).

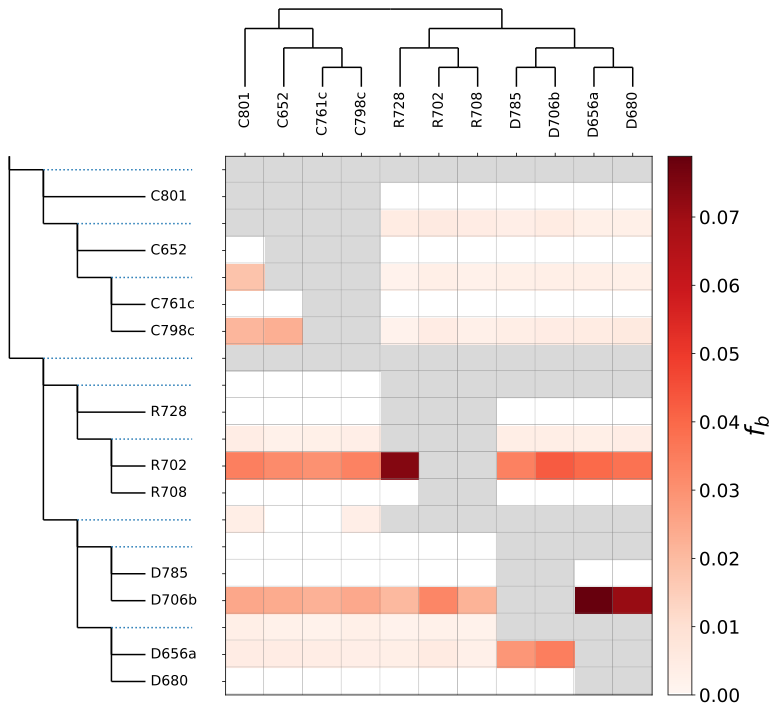

**Figure S7. F-branch estimates demonstrate widespread introgression.** F-branch [$f_{b}(C)$] statistics were estimated using a RAxML-inferred phylogeny (Fig. 1D). F-branch statistics (red hue scale) are a summary of f4-admixture ratio scores, which capture excess allele-sharing between a branch $b$ (y-axis) and population or individual $C$ (x-axis) relative to the sister lineage of branch $b$. *P. pilosa* was used as the outgroup for all comparisons. Individual tips are labeled with the first letter of the individual’s species identity (e.g., “D” for *P. drummondii*).

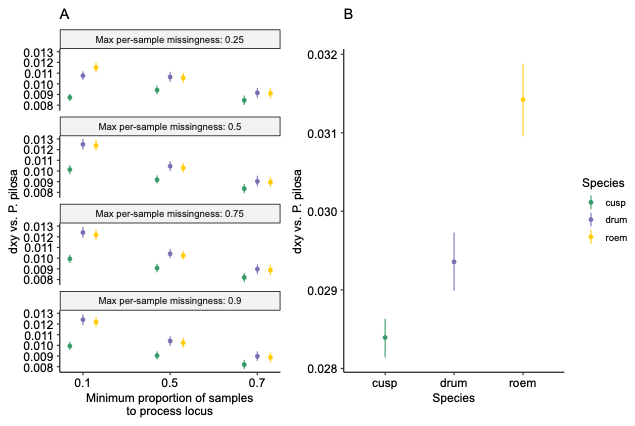

**Figure S8. Genetic divergence (**$\boldsymbol{d}_{\boldsymbol{xy}}$**) across datasets.** Genetic divergence ($d_{xy}$) estimates using reduced representation sequencing (RRS, a) and whole genome sequencing datasets (WGS, b) between outgroup *P. pilosa* and annual species *P. cuspidata*, *P. drummondii*, and *P. roemeriana*, drawn in green, purple, and red, respectively. RRS estimates are presented with respect to multiple filtering criteria. The x-axis corresponds to the minimum proportion of samples in which a RAD-Seq locus must be present to be retained (equivalent to the -r and -R flags in the stacks-populations tool) and panels correspond separate estimates by the maximum missingness rate required to retain an individual sample (equivalent to the -mind flag in plink v1.9).

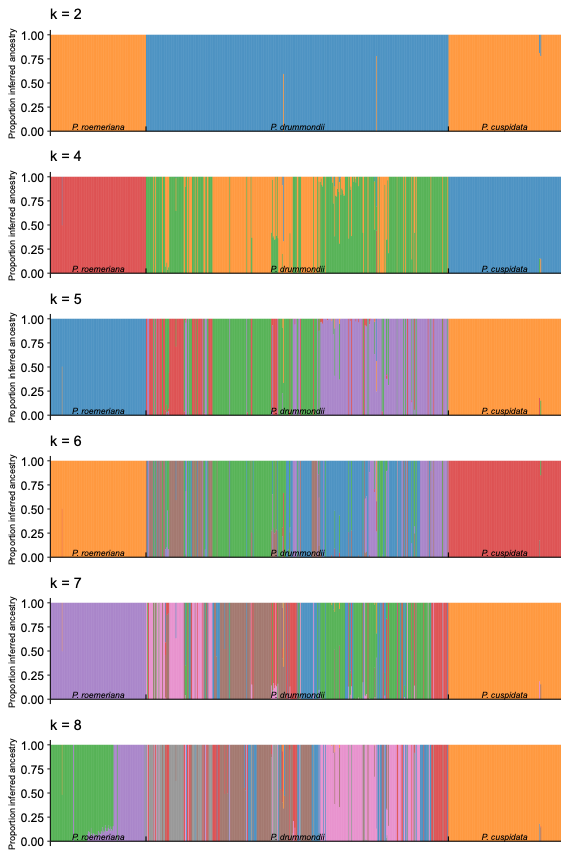

**Figure S9. ADMIXTURE plots.** Mixing proportions for $K=2, 4, 5, 6, 7, 8$ ancestral populations determined with ADMIXTURE using the RRS dataset as in Fig. 2B. Heights of colored bars denote estimated ancestry proportions corresponding to each of the K populations. Individuals are grouped by species label (black ticks) and within each group arrayed from left to right by increasing longitude.
